# NeuroAtlas: An Artificial Intelligence-based Framework for Annotation, Segmentation and Registration of Large Scale Biomedical Imaging Data

**DOI:** 10.1101/2024.08.24.609507

**Authors:** Hassan Mahmood, Farah Nawar, Syed Mohammed Shamsul Islam, Asim Iqbal

## Abstract

With increasing neuroimaging modalities and data diversity, mapping brain regions to a standard atlas template has become a challenging problem. Machine learning in general and deep learning, in particular, have been providing robust solutions for several neuroimaging tasks, including brain image registration and segmentation. However, these methods require a large amount of data for groundtruth labels, annotated by human experts, which is time-consuming. In this work, we introduce NeuroAtlas, an AI-based framework for atlas generation and brain region segmentation. We showcase an end-to-end solution for brain registration and segmentation by providing i) a deep learning modeling suite with a variety of high-performing model architectures to map a brain atlas onto the input brain section and ii) a Graphical User Interface (GUI)-based plugin for large-scale data annotation with a feature of modifying the predicted labels for active learning. We demonstrate a robust performance of our framework on the human brains, captured through various imaging modalities and age groups, and demonstrate its application for mouse brains as well. NeuroAtlas tool will be open-sourced and entirely compatible with both local as well as cloud-based computing so that users can easily adapt to their neuroimaging custom datasets.

## 1. Introduction

Understanding the brain has been a subject of mystery for a long time. In this pursuit, neuroimaging technologies are developed to capture the anatomical structures and functional principles in the brain. To analyze such highly complex and large imaging datasets, the limiting accessibility of the right tools and technologies has been one of the detrimental factors. With the rapid growth in Artificial Intelligence (AI), the availability of data annotation tools and the performance of image registration and segmentation methods have been notably enhanced for Computer Vision (CV). This, however, is not translated in neuroimaging, at its full capacity, due to the challenges in adaptability towards CV-based technologies.

Through advances in neuroimaging techniques, there exists an inherent challenge to analyze the data due to the complexity of brain imaging datasets and the subsequent challenges towards large-scale data annotation. Henceforth, mapping the brain sections to a common atlas through image registration or segmentation remains an existing challenge. This issue can be addressed by bridging the gap between AI-based techniques and neuroimaging. Recent efforts in this direction have resulted in the development of data annotation tools to assist in generating ground-truth labels for selected images v7labs [5], labelbox [2], SuperAnnotate [4], Dataloop [1], Segmentai [3] few to mention, some are mentioned in Table 1. These tools are well established. Some of them are best for user-friendliness (like Labelbox or SuperAnnotate), and some are known for their cutting-edge automation (V7). However, these have price tags on them. There exist some open-source tools for medical image segmentation with some pros and cons to them. Dicomworks [21] is a good tool for anonymization of the data, annotation, and measurements but lacks in creating binary masks for the user. ITK-Snap [23] is quite flexible software for data labeling. This tool provides a complete 3D view during annotation but lacks in providing bounding box annotation. It worked well with NiFTI [13], NRRD, and VTK Image formats. LabelMe [19] is a lightweight user-friendly data annotation with the ability to export annotated images into different popular formats like the COCO format, and the Pascal-VOC format. However, it provides limited scalability and is not suitable for working with complex content. RIL-contour [20] and 3D-Slicer [14] are well-established free tools, however, these are quite extensive, and the learning curve is too large to be able to efficiently use these tools.

**Table 1.** List of Free and Paid Annotation Tools.

| Name | Price |
| --- | --- |
| Dicomworks | Free |
| ITK-Snap | Free |
| 3D-Slicer | Free |
| RIL-contour | Free |
| LabelMe | Free |
| v7labs | Paid |
| Labelbox | Paid |
| SuperAnnotate | Paid |
| Dataloop | Paid |
| Segmentai | Paid |

In terms of AI-based frameworks like SegmentAi [3], most of the existing efforts are providing automated segmentation which users can correct later, but most of them are not freely available and also extensive in learning, causing hesitation for the researchers to adopt those tools.

Besides that, there exists a limitation in the existing solutions, for instance, most data annotation tools are not designed to take the labeling format (e.g. binary masks and polygons) into account to facilitate the AI-based neuroimaging analysis pipeline and also the availability of some of the solutions are not straightforward, some of them are costly while others are extensive. Moreover, despite the advancements in cloud computing technology most freely available tools are still based on local machine installation. Similarly, the existing ML-based solutions provide a single model (registration or segmentation algorithm), designed for a particular dataset or image modality. This hinders the generalisability of the ML-based solution towards its adaptability to different datasets in the non-computer science community.

Ideally, the goal is to design a single solution that provides a multi-modal framework with state-of-the-art image registration and segmentation architectures, adapted for a variety of neuroimaging datasets. Moreover, since the DNN-based techniques are data-hungry, providing a feature of ‘active learning’in a single framework will be very useful for the neuroscience community. In this way, single or multiple users can start with minimum annotated samples, train the model on the labeled data, and use this trained model to smartly assist in generating the ground-truth labels for the remaining training set. In addition, once the annotation process is completed, selecting the optimal model is a tedious task, partly because each network is designed for a unique dataset or a different task and partly due to a lack of computing knowledge base among the medical community. A unified framework that guides the user toward selecting an optimal model from a model zoo will be of huge assistance in designing a large-scale AI-based neuroimaging analysis pipeline.

In this study, we introduce NeuroAtlas, an AI-based unified framework to generate labels through a GUI-based interactive active learning process and to apply state-of-the-art deep learning models. NeuroAtlas is designed to take a high-dimensional training dataset as input, generate the embeddings through Variational AutoEncoders (VAEs) [9] in a low dimension, and then apply a clustering algorithm to discover the minimum samples needed for labeling to cover the variational space in the entire dataset. This is to assist the user in generating the minimum ground-truth labels for training the model towards active learning. This will not only result in saving a huge amount of time for data annotation but will also pave the way towards few-shot learning i.e. minimum labels needed to achieve decent performance on a novel dataset or task.

Moreover, our framework is designed to automatically convert the ground-truth labels into the AI models’ compatible format as well as guide the user in selecting the optimal model for the task at hand. NeuroAtlas is compatible with both 2D images and 3D volumes, and we demonstrate its performance on a variety of registration and segmentation tasks for human, mouse, and drosophila-based neuroimaging datasets. We provide the framework to smoothly run in both local as well as server-based computing hardware setup and provide a model-zoo of trained deep neural networks where the users can select them for their similar datasets as shown in Fig. 1. We believe NeuroAtlas will serve as a single hub for the neuroimaging community to apply and extend the AI-based registration and segmentation models to cover all kinds of biological datasets in the future.

**Figure 1.**
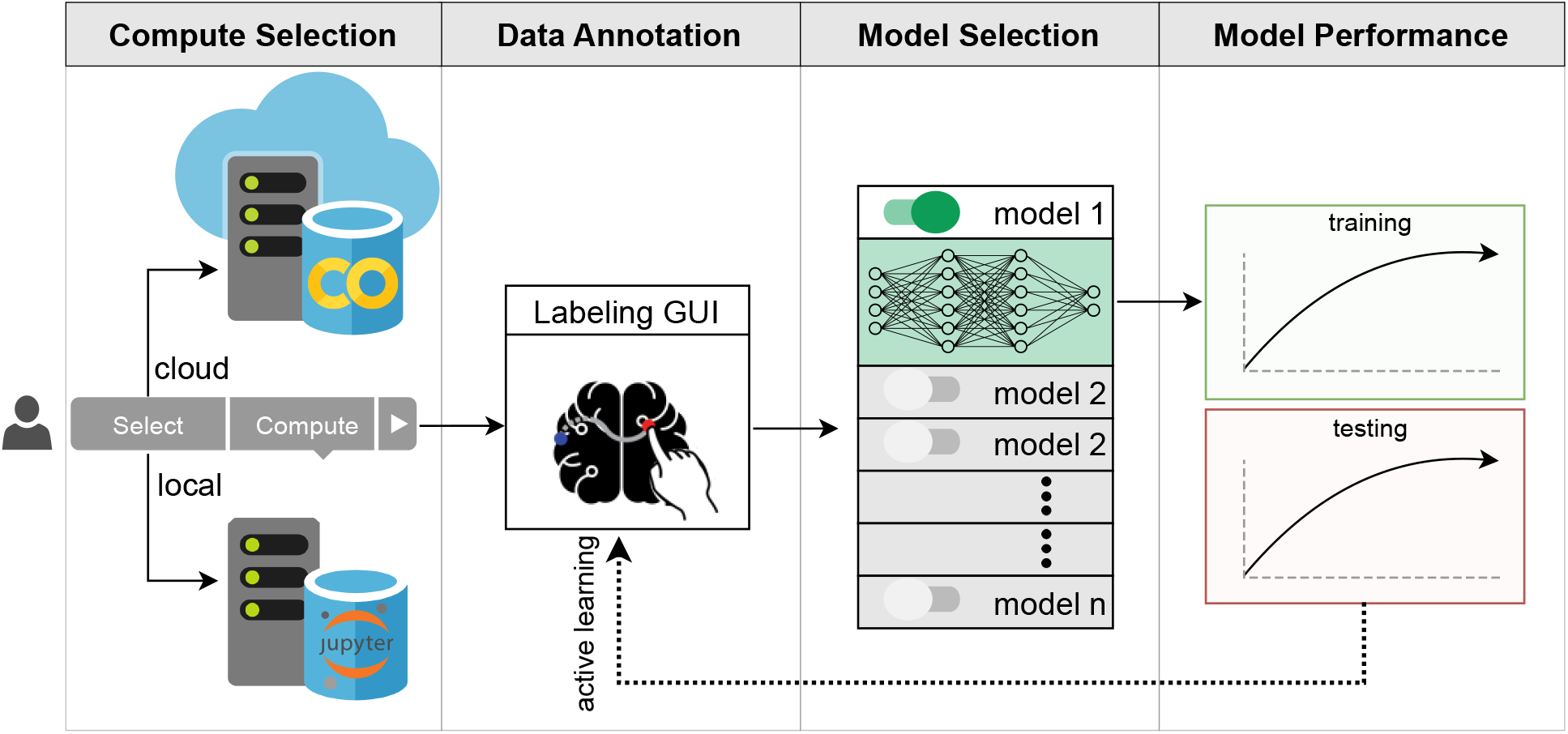
NeuroAtlas block diagram: User can select the compute type to run locally or through Google Colab. Once the compute is selected, the annotation GUI supports the user to load and annotate the labels on the selected training set. After ground-truth data is generated, the user can select the corresponding segmentation or registration model to run training. Once the model is trained, the user can load a pre-trained model to help in future data annotation through active learning.

## 2. Methods

In this section, we explain the models and functionality provided by our framework. A guideline of how and when to use specific models/methods is also described.

### 2.1 Annotation

Fig. 2 illustrates the steps for using our annotation tool. First, a user needs to provide data and label paths and also name and number of regions they want to annotate. By selecting the desired region name from the drop list, the user can annotate as many regions as required using the freeform tooltip in the app. In step 4 of Fig. 2, the corresponding binary masks of slice number 15 written in storage space are shown. After annotating the shortlisted dataset from the clustering curation step, it can be used to train the model and get new segmentation results on the test data. The sample output from 2D U-Net can be seen in Fig. 3. We can correct these masks using our annotation app. Fig. 4 demonstrates the usage of our tool for correcting the labels, which can be further added into the training of the U-Net Model afterward.

**Figure 2.**
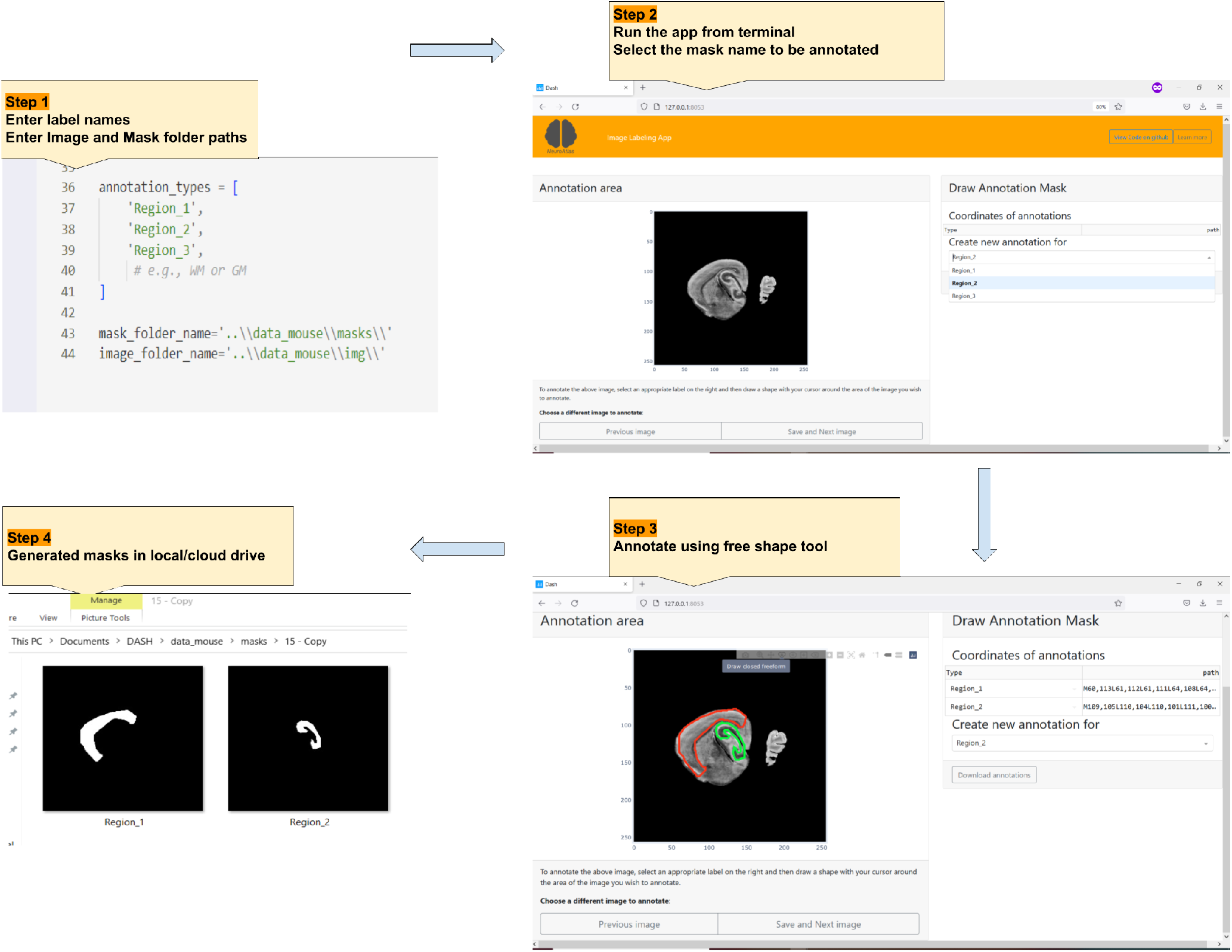
Annotation GUI: Step1-4 describes the flow for new data annotation on a sample image taken from the mouse brain dataset. Users can select the label type and start drawing the ground-truth mask through interactive polygons. The progress is saved throughout the session.

**Figure 3.**
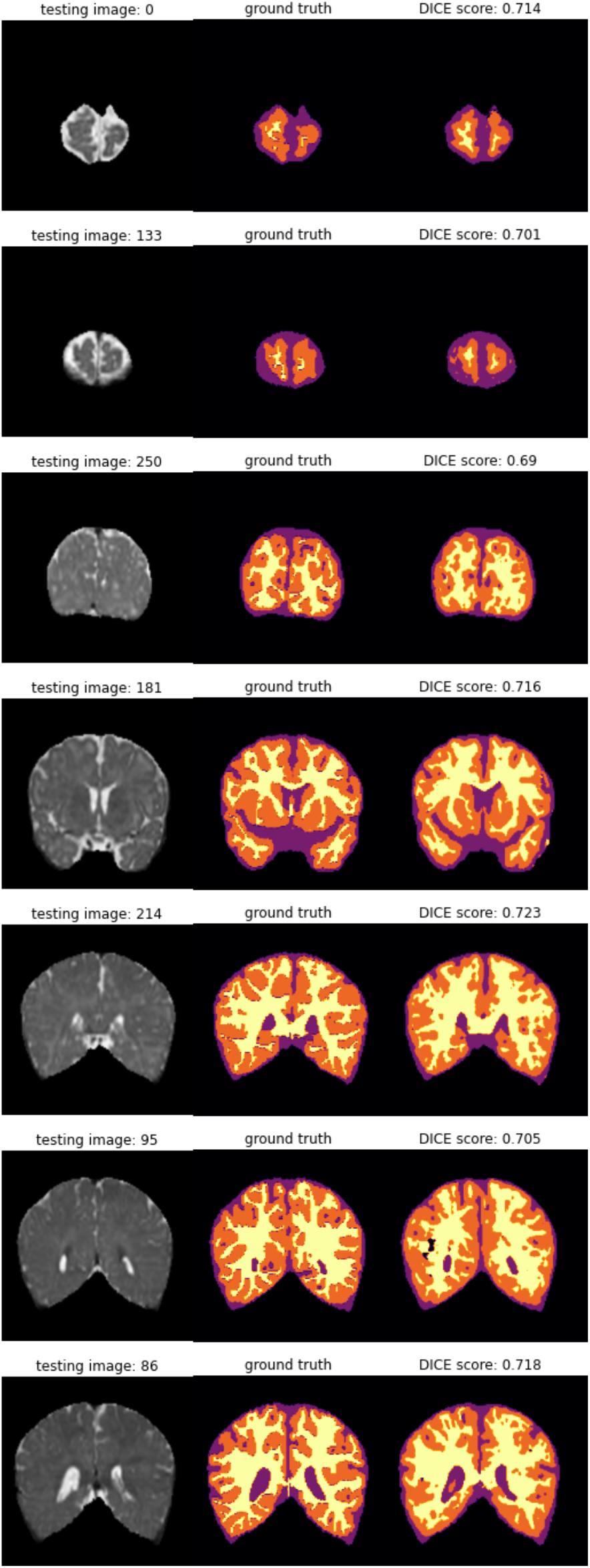
Sample segmentation output of the 2D U-Net is demon-strated which is trained to segment human brain MRI containing White Matter (WM), Grey Matter (GM), and Cerebrospinal Fluid (CSF).

**Figure 4.**
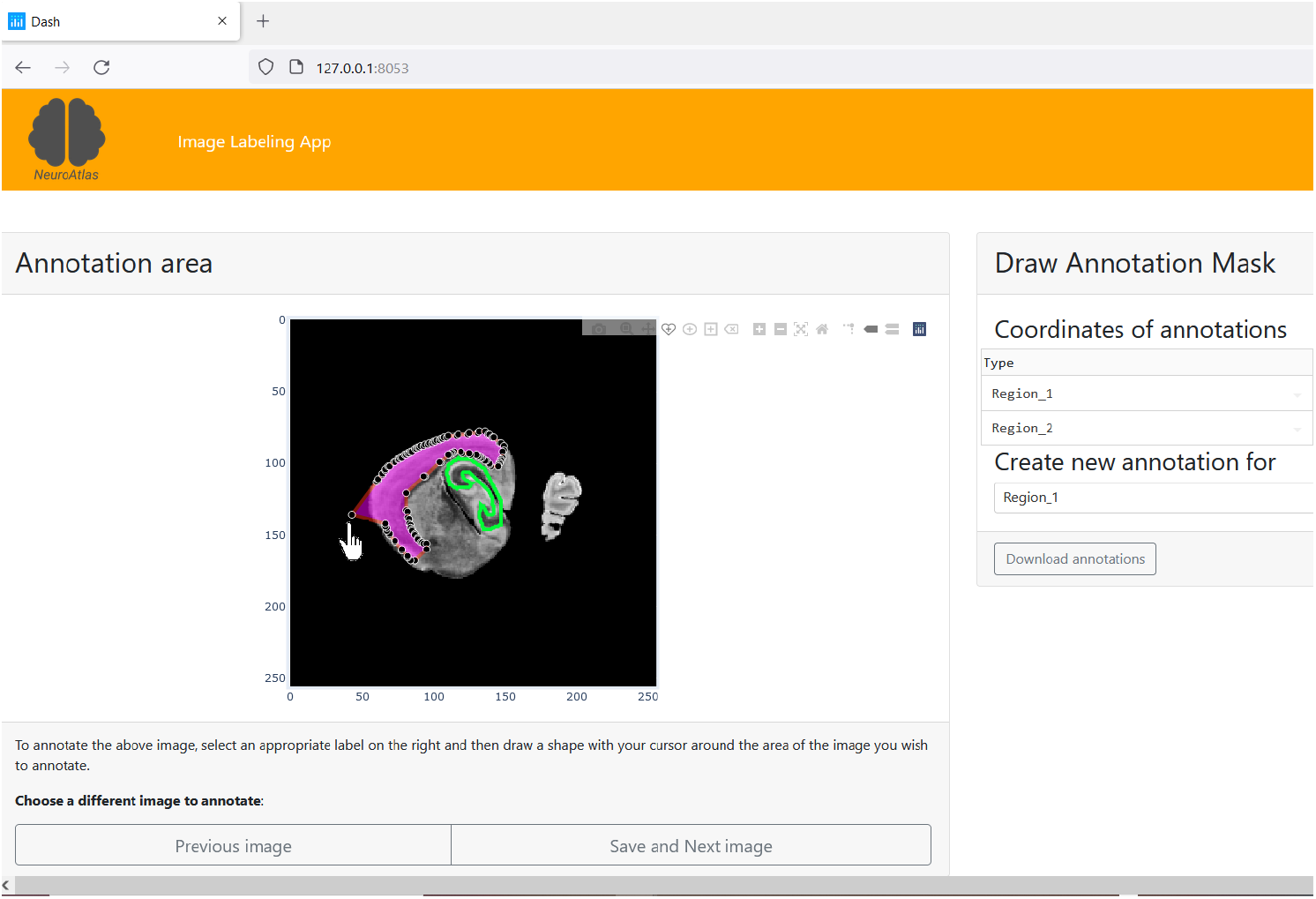
Demonstration of correcting the already assigned/predicted labels using annotation app

### 2.2 Image Registration

The purpose of the image registration task is to find the mapping between two images. The transformational model for mapping can be rigid, affine, or non-rigid, depending on the nature of the task. Image registration is a useful tool to conduct longitudinal studies of different biological or psychophysical phenomena. This can also be used for atlas-based segmentation. The performance of atlas-based segmentation largely depends on the quality of registration and fusion methods for multi-atlas-based segmentation. Some of the famous registration tools provided by our framework are explained below.

#### 2.2.1 Advanced Normalization Tools (ANTs)

The ANTs [6] open-source software library consists of a suite of image registration and template-building tools for quantitative morphometric analysis. This tool provides a lot of flexibility by giving different options to the user. This library is mainly based on classical image registration algorithms. The usual flow of preprocessing in registration is to first rigidly align the images, then depending upon the nature of the data affine registration is performed. Finally, non-rigid registration is applied to get the change in the region of interest. This library is available in C, Python, and R versions. Complete information about the practical examples can be found in ANTs-Tutorial.

For rigid or affine transformation in ANTsPy (Advanced Normalization Tools in Python), the syntax is used as shown below

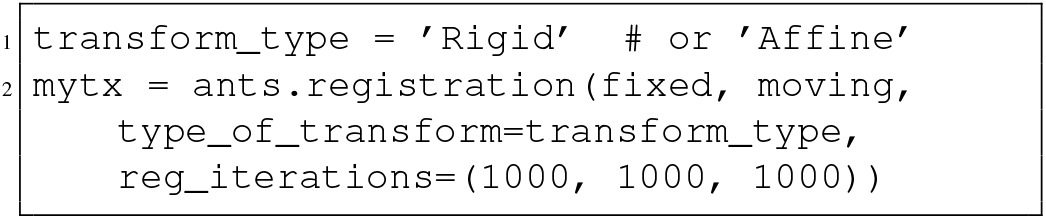

Here, reg iterations define the number of iterations at three scales. Afterwards, non-rigid registration can be applied as following settings:

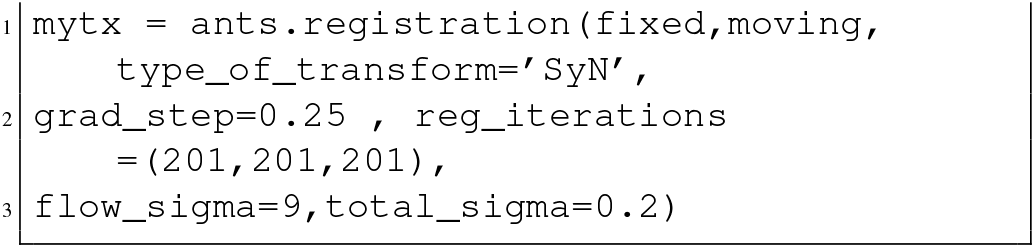

Transformations from each step can be applied on other images or labels as below:

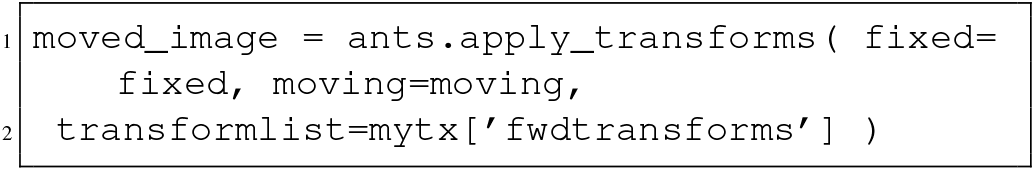

#### 2.2.2 Deep Learning-based Registration Models

Learning-based registration models provide comparable performance with excellent time efficiency. These models would be useful to conduct population studies in general. We include VoxelMorph [7] and DeepIntense [17, 18] in our framework.

**VoxelMorph** is one of the deep learning-based image registration methods. This architecture is based on a U-Net [22], spatial transformer network (STN) [12], and a regression layer for generating a deformation field. The model is trained using an MSE-based loss function using Adam optimizer. During inference, the model’s output is the warped image and flow field. The flow field is later used to deform other images or labels.

**DeepIntense** work is inspired by VoxelMorph. This method proposes a new loss function to be used in the image registration deep learning architecture. The prime focus of this work is to reduce the effect of the change in intensity in the data using perceptual loss during inference.

For a patient-specific and critical task like surgery planning, classical registration methods would be preferable as they provide more theoretical guarantees. Learning-based models are lacking in providing theoretical guarantees like diffeomorphism [8] and smoothness of the deformation field at large. Diffeomorphism is about the invertibility of the deformation field where smoothness is required to stop any unnatural transformation. There are some efforts to achieve diffeomorphism in learning-based models [8], but they are lacking in general. These learning-based models are mainly for non-rigid registration. Rigid or affine transformation needs to be performed beforehand.

### 2.3 Image Segmentation

#### 2.3.1 U-Net

U-Net is an encoder-decoder-based semantic segmentation architecture. It has skip connections between encoding and decoding layers to provide semantic context for the segmentation prediction. For our implementation, the U-Net input layer size is kept at 256x256. We have trained the network with a bottleneck size of 2048 using DICE-based loss and an Adam optimizer by keeping a learning rate of 0.0001. After training the model on the training dataset, it has been validated on the testing data. We provide both 2D and 3D implementation of U-Net architecture with an option to customize the input size and depth of encoder-decoder architectures.

#### 2.3.2 SeBRe: Segmenting Brain Regions

SeBRe is an instance segmenter with Mask RCNN as the background architecture. Mask RCNN [10] is an extended version of Faster R-CNN such that it predicts an object mask alongside object detection. Mask R CNN performs instance segmentation i.e. it can detect and segment different instances of the same class. SeBRe [11] is an extended version of Mask R-CNN, which is constructed by using the first five blocks of ResNet101 and Feature Pyramid Network (FPN). The network architecture is the same as of [10]. Region Proposal Network (RPN) processes the input feature map in a sliding-window fashion which forwards the predicted n Regions of Interest (ROI) from each window to the Mask R-CNN ‘heads’ based on the Fully Pyramidal Network (FPN). In multi-headed FPN, the classifier head acts as an identifier, the regressor head detects the regions and predicts bounding boxes around the ROI, while the segmenter predicts the mask using Fully Convolutional Network (FCN) [16]. We apply transfer learning by using pre-trained weights of the Microsoft COCO dataset [15].

## 3 Conclusion

In this manuscript, we introduce NeuroAtlas, a machine learning-based framework to assist in smart data annotation, segmentation, and registration tasks in both 2D and 3D imaging modalities. We use VAEs to explore the minimum data samples needed to represent maximum data variability in training. This will provide ease for biologists and neuroscientists while decision-making about shortlisting their training and testing data. Furthermore, our annotation tool in the NeuroAtlas framework provides an interactive platform to draw, annotate, and modify the ground truth and predicted labels. In addition, we introduce an active learning-based feature to minimize the workload in generating ground truth data from the existing data modality. We also provide ready-to-use implementations of a variety of deep learning models such as U-Net (2D and 3D), SeBRe (Mask R-CNN), VoxelMoprh (2D and 3D), and DeepIntense (2D and 3D). To facilitate the community, we open-source the aforementioned models trained on publicly available datasets using human and mouse databases. Users can adapt and improve the existing models as well as train them from scratch on their custom datasets. With the support of the community, we aim to extend our model zoo in the future to cover both micro and macro imaging samples from the existing and new data modalities. We believe NeuroAtlas will serve as a single platform to facilitate the users in running large-scale data analytics using our end-to-end deep learning-based framework.

